# Genetic Identification of Novel Separase regulators in *Caenorhabditis elegans*

**DOI:** 10.1101/191452

**Authors:** Michael Melesse, Dillon E. Sloan, Joseph T. Benthal, Quincey Caylor, Krishen Gosine, Xiaofei Bai, Joshua N. Bembenek

**Affiliations:** Department of Biochemistry, Cellular and Molecular Biology, University of Tennessee, Knoxville, TN

**Keywords:** Separase, PPH-5, HSP-90, Suppressors, C. elegans

## Abstract

Separase is a highly conserved protease required for chromosome segregation. Although observations that separase also regulates membrane trafficking events have been made, it is still not clear how separase achieves this function. Here we present an extensive ENU mutagenesis suppressor screen aimed at identifying suppressors of *sep-1(e2406)*, a temperature sensitive maternal effect embryonic lethal separase mutant. We screened nearly a million haploid genomes, and isolated sixty-eight suppressed lines. We identified fourteen independent intragenic *sep-1(e2406)* suppressed lines. These intragenic alleles map to seven SEP-1 residues within the N-terminus, compensating for the original mutation within the poorly conserved N-terminal domain. Interestingly, 47 of the suppressed lines have novel mutations throughout the entire coding region of the *pph-5* phosphatase, indicating that this is an important regulator of separase. We also found that a mutation near the MEEVD motif of HSP-90, which binds and activates PPH-5, also rescues *sep-1(e2406)* mutants. Finally, we identified six potentially novel suppressor lines that fall into five complementation groups. These new alleles provide the opportunity to more exhaustively investigate the regulation and function of separase.

## INTRODUCTION

Separase is a highly conserved cysteine protease required for proper chromosome segregation during anaphase of both meiotic and mitotic stages of cell division (Peters et al. 2008). Separase proteolytic activity is inhibited during interphase and early mitosis by its pseudosubstrate inhibitor, securin (Nasmyth, K. A., 2002). The protease activity of separase is critical for the cleavage of kleisin subunits of the cohesin complex (Uhlmann et al. 2000, Hauf et al. 2001). Cohesin holds sister chromatids together prior to their proper attachment to spindles and alignment on the metaphase plate preceding anaphase (Nasmyth and Haering, 2009). Separase has also been implicated in various cell cycle regulatory functions. In budding yeast, separase stabilizes the anaphase spindle by cleaving the spindle and kinetochore associated protein, Slk19 (Sullivan et al. 2001). It is also involved in the release of the essential mitotic phosphatase Cdc14 in budding yeast (Sullivan and Uhlmann 2003). In mammalian cells, separase licenses centriole duplication (Baskerville et al. 2008) and a domain within its N-terminus binds and inhibits the Cyclin B-Cdk1 complex (Gorr et al. 2005). In mammalian cells, separase has also been shown to associate with membranes and its depletion is associated with swelling of the trans-golgi network and decreased constitutive protein secretion (Bacac et al. 2011). In *Arabidopsis*, separase mutant cells display mitotic failure due to defects in vesicle trafficking along microtubules, which is critical for synthesis of a cell plate during cytokinesis (Moschou et al. 2016). Therefore, there are numerous functions of separase during the cell cycle, and how each are regulated has not been fully elucidated.

In *Caenorhabditis elegans*, separase is known to regulate multiple cell cycle events beyond its chromosome segregation functions (Bembenek et al. 2007, 2010). It has been demonstrated to regulate cell cycle related membrane transport events critical for both cytokinesis and embryonic development. During meiosis, the *C. elegans* eggshell is formed around a fertilized embryo to prevent polyspermy and provide both mechanical as well as osmotic protection for the developing embryo (Olson et al. 2012; Stein and Golden 2015). Formation of the eggshell is dependent on the progression of the embryonic cell cycle and requires cargo released via cortical granule exocytosis (CGE), which occurs during anaphase I (Bembenek et al. 2007). Importantly, separase localizes to cortical granules and is required for their exocytosis during anaphase, independently of chromosome segregation.

Various separase mutants have been identified in budding yeast, mouse and human cells. Many of these mutants compromise the protease function of separase and directly affect its role during chromosome segregation. Interestingly, the hypomorphic separase mutant (*sep-1(e2406)*), originally isolated by David Livingstone in a screen for temperature sensitive mutants defective in cell division (Siomos et al. 2001), is a partial separation of function allele. *sep-1(e2406)* is a C450Y missense mutation in the N-terminal region of separase and has minimal effect on the chromosomal segregation role of separase but significantly diminishes cortical granule exocytosis. In embryos, SEP-1(e2406) can be observed on the spindle, but shows reduced localization to cortical granules and results in a lower number of exocytic events. Another separase mutant (*sep-1(ax110)*) is a non-conditional allele that also results in minimal chromosome segregation defects and leads to cytokinesis failure (Richie et al. 2011). This allele is a missense mutation (H738P) in the protease domain of SEP-1 that is maternal effect embryonic lethal. These alleles potentially provide a unique opportunity to learn more about the membrane trafficking functions of separase.

Previous attempts to learn more about separase regulation used *sep-1(e2406)* to identify the PPH-5 phosphatase as a suppressor of separase (Richie et al. 2011). This effort screened 1.0 X 10^5^ genomes and identified three suppressors including one *pph-5* allele, *pph-5(av101)*, an intragenic *sep-1* (L556F) mutant and another mutant that maps to LG III. Mutations in *pph-5* as well as RNAi mediated knockdown rescues *sep-1(e2406)* (Richie et al. 2011), suggesting that *pph-5* is a negative regulator of separase function. The *pph-5(av101)* suppressor allele, which is a missense mutation (P375Q), does not suppress *sep-1(e2406)* at 24° but was effective in suppressing *sep-1(ax110)* at all tested temperatures (Richie et al. 2011). This observation suggests that there might be underlying differences in the effects of these SEP-1 mutations on separase function.

PPH-5 is a widely conserved phosphatase that contains N-terminal tetratricopeptide repeats (TPRs) and a C-terminal phosphatase domain. PP5 (human PPH-5), originally identified as a regulator of a variety of cellular signaling pathways including glucocorticoid receptor signaling, displays low phosphatase activity when purified due to the autoinhibitory role of its TPR domain (Chen et al. 1996). Interestingly, PP5 binds CDC16 and CDC27, components of the Anaphase Promoting Complex/Cyclosome (APC/C) (Ollendorff and Donoghue 1997). The APC/C is an E3 ubiquitin ligase required for activation of separase at the metaphase to anaphase transition and is regulated by phosphorylation (Kraft et al. 2003; Chang and Barford 2014; Musacchio 2015). The precise mechanism by which PPH-5 regulates separase is unknown, but these findings suggest that it may be an important regulator of the metaphase to anaphase transition.

One of the well-studied regulatory pathways of PPH-5 is its interaction with the molecular chaperone HSP-90. The crystal structure of auto-inhibited human phosphatase 5 (PP5) shows that access to the enzyme active site is blocked by a combination of the TPR domain and a C-terminal *a*J-helix (Yang et al. 2005). HSP-90 binds the TPR domain of PPH-5 to release auto-inhibition and promote phosphatase activity towards protein substrates (Haslbeck et al. 2015). HSP-90 consists of three highly conserved domains and binds its client proteins via its middle domain (MD), while it binds co-chaperones via its C-terminal domain (Schopf et al. 2017). The very C-terminal MEEVD motif is critical for HSP90 interaction with TPR domain containing co-chaperones like PP5. As a major protein chaperone, HSP-90 is known to bind multiple proteins (Haslbeck et al. 2013). Available HSP-90 mutants as well as RNAi in *C. elegans* cause penetrant pleiotropic phenotypes (Inoue et al. 2006; Gillan et al. 2009; Gaiser et al. 2011). To our knowledge, there is no evidence linking HSP-90 to regulation of separase in any system.

In this paper, we present the results of a genetic suppressor screen aimed at uncovering regulators of separase. We identified intragenic suppressors, *pph-5* mutants, a novel *hsp-90* allele and unknown alleles that fall into five complementation groups. These suppressors may provide important insight into separase regulation and function.

## MATERIALS AND METHODS

### Mutagenesis and Selection

Strains were maintained as described (Brenner 1974). *sep-1(ax110)* screen: *sep-1(ax110)*/hT2 [*bli-4(e937) let-?(q782) qIs48 (Pmyo-2::gfp; Ppes-10:: gfp; Pges-1::gfp)*] (I,III) worms were synchronized by bleaching with hypochlorite and grown to L4. Mutagenesis was performed by incubating worms with 0.5mM ENU for 4 hours at 25° and recovering in 50ml of M9 overnight at 15°. 30 P_0_s were plated to 81 100mm plates, transferred to 25° and incubated. After one generation, 50 unbalanced (non-green, should be homozygous for *sep-1(ax110)*) F_2_ progeny from each of the 81 100mm plates were moved onto 60mm OP50 plates and checked for fertility. From each non-green F_3_ producing plate, at least 6 plates of non-green animals were cloned and genotyped. Candidate suppressed lines were confirmed to be homozygous for *sep-1(ax110)* and sequenced for mutations at the *pph-5* locus.

*sep-1(e2406)* screen: homozygous *sep-1(e2406)* worms were synchronized by bleaching with hypochlorite and grown to L4. Worms were mutagenized with 0.5mM ENU in M9 for 4 hours and recovered in 50ml of M9 for 1 hour at 15°. 100 mutagenized worms were moved to each of 60 MYOB plates and incubated at 15°. P_0_s were moved to new plates daily. The number of F_1_ worms on each plate were estimated and plates were grown for multiple generations at 15°. These plates were then chunked and incubated at 20° and allowed to produce offspring. Plates that yielded embryos were cloned and backcrossed to *sep-1(e2406)* for multiple generations.

### Identification of suppressor mutations

Genotyping: *sep-1(ax110);* primers (oASP-UTK-3 and oASP-UTK-4) were used to amplify a *sep-1* fragment by PCR. The PCR product was then digested with a restriction enzyme (SacII), which is introduced by the *sep-1(ax110)* mutation. The *sep-1(e2406)* allele was genotyped by sequencing a PCR fragment amplified using a pair of primers (oASP-UTK-34 and oASP-UTK-29) and sequenced with oASP-UTK-7.

PCR and sequencing: PCR primers were used to amplify the locus of interest from worm lysates. PCR products were then gel purified and sequenced. Three PCR fragments of *sep-1*, five of *pph-5* and two of *hsp-90* were amplified, spanning across each gene. Primers used for PCR and sanger sequencing of *sep-1*, *pph-5* and *hsp-90* loci are listed in supplementary tables (Tables S2, S3 and S4).

### Characterization of suppressed lines

Hatching assay: four P_0_ L4 larvae were placed in each of 35mm OP50 NGM plates and allowed to lay embryos for 24 hours at the experimental temperature (15°, 20° or room temperature). Worms were then moved to new plates and returned to temperature to continue laying embryos. The number of embryos and hatched animals on overnight plates was counted on each plate and plates were incubated for 24 hours. The following day, the number of unhatched embryos or hatched larvae was counted and % hatching was quantified.

RNAi feeding: Worms were moved onto NGM plates with ampicillin and isopropyl-b- D-thiogalactopyranoside which were seeded with HT115(DE3) bacteria carrying RNAi feeding constructs for 24 hours. Worms were then moved onto new RNAi feeding plates daily and hatching embryos were counted. Unless otherwise stated, five L1 stage worms per strain were fed at 20°. Animals were moved to new RNAi feeding plates after reaching the L4 stage, and hatching was quantified daily for 48 hours.

Western blot analysis: Worms were grown at 20° on 100mm OP50 seeded plates for one generation and collected by washing in M9 buffer. Each worm pellet was resuspended in 1xSDS loading buffer (2μl/mg of pellet) and heated in a microwave (4x20 sec with 1 min cooling). Lysates were then centrifuged (15,000 x RCF, 10 min) and supernatant was transferred into new tubes. 10 *μ*l of worm lysate was then loaded per well and analyzed by standard western blot. SEP-1 was detected by using a polyclonal rabbit antibody (Richie et al. 2011) at a dilution of 1:750. Secondary antibody used was anti-rabbit 700 from Li-Cor and quantified using the Image Studio software. Non-specific bands were used to normalize signals between lanes. All antibody incubations were done in the presence of 5% (w/v) non-fat milk.

Complementation tests: Twenty-five males generated using *him-5 RNAi* bacterial feeding for each strain were mated with five hermaphrodites on unseeded NGM plates and incubated at 20° for 24 hours. Mated worms were then moved to OP50 seeded 60mm plates and allowed to lay F_1_ embryos at 15°. Once F_1_ worms reach L4 stage and the presence of ∼50% male animals was observed, indicative of successful mating, four L4 hermaphrodites in triplicate were moved to OP50 seeded 35mm NGM plates and incubated at 20°. Viability of F_2_ embryos was determined.

### Reagent Availability

All strains are available upon request.

## RESULTS AND DISCUSSION

### Identification of suppressors

To identify genes that regulate separase function, we performed mutagenesis screens for suppressors of two separase mutants, *sep-1(ax110)* and *sep-1(e2406)* (Figure 1A). We first screened for suppressors of the non-conditional *sep-1(ax110)* mutant (Figure S1A) which introduces a point mutation in the protease domain of SEP-1(H738P) and is maternal effect embryonic lethal. We postulated that this separase allele might be differentially impaired relative to the temperature sensitive *sep-1(e2406)* allele, which introduces a mutation in the TPR-like domain (C450Y) and might be suppressed by a different set of mutations. This suppressor screen identified four independent suppressors of *sep-1(ax110)*, all of which were *pph-5* mutants (*erb1(S229L)*, *erb2(M380T), erb3(L77P)* and *erb4(L77P)*) from 56,404 genomes screened (Figure 1B and Figure S1B). This is consistent with a previous finding that *sep-1(ax110)* is completely rescued by loss of *pph-5* (Richie et al. 2011). Therefore, we focused our efforts towards identifying suppressors of *sep-1(e2406)*.

**Figure 1.**
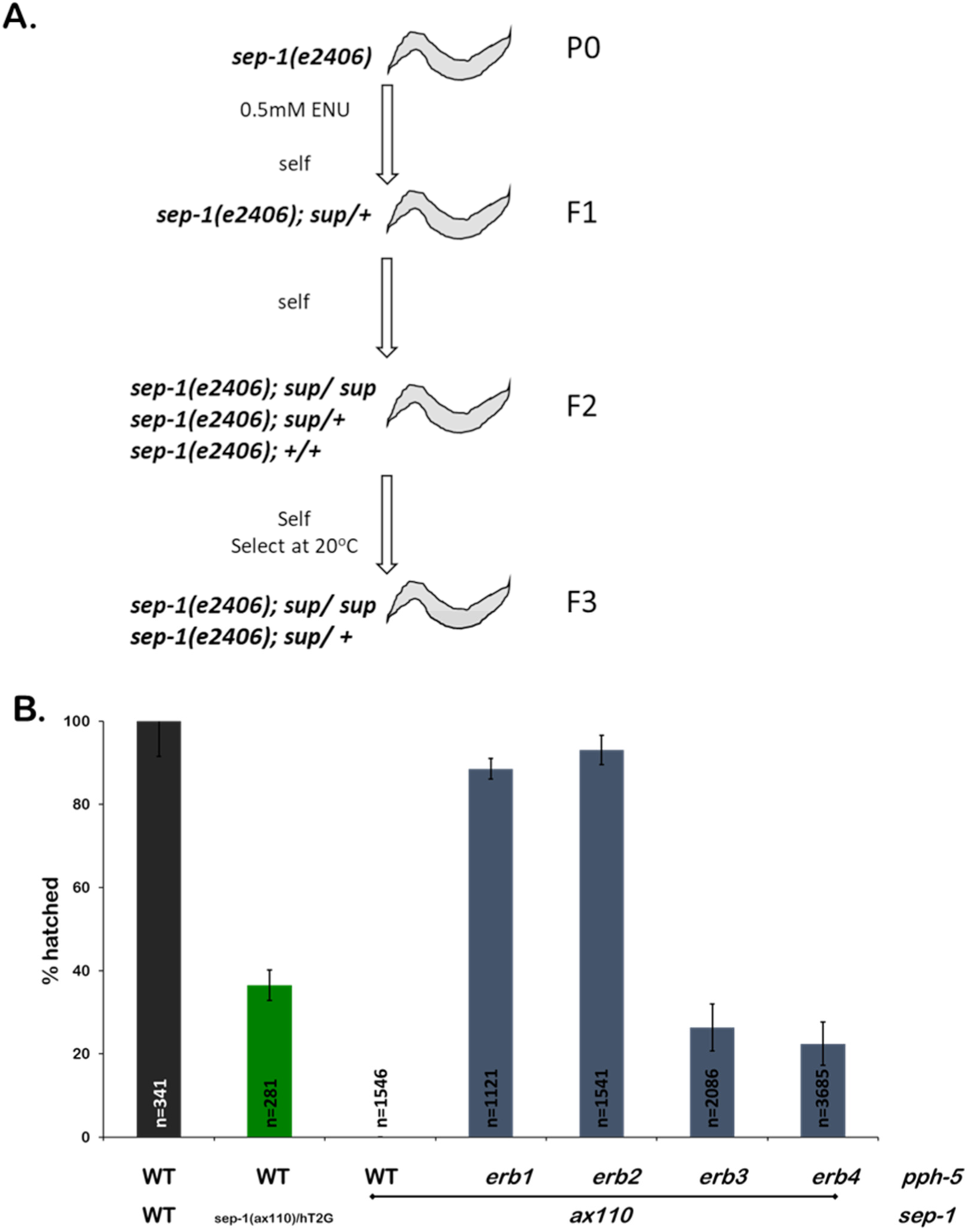
Isolation of suppressors of *sep-1(ax110)* and *sep-1(e2406)* **A.** Schematic for the isolation of lethality suppressing mutants in the temperature sensitive *sep-1(e2406)* background via ENU mutagenesis. **B.** Mutations in *pph-5* rescue non-conditional *sep-1(ax110)* mutants. *sep-1(ax110)* homozygotes carrying mutations in the phosphatase domain, *erb1* (S229L) and *erb2* (M380T), of PPH-5 have lower embryonic lethality relative to those carrying mutations in the TPR domain *erb3/4* (L77P) (n=number of embryos).

The *sep-1(e2406)* mutation results in a temperature sensitive maternal effect embryonic lethality. When L4 animals are shifted to 20°, the lowest temperature at which lethality is fully penetrant, *sep-1(e2406)* hermaphrodites lay 100% dead embryos. *sep-1(e2406)* embryos are unable to perform cortical granule exocytosis and fail to build an eggshell when maintained at 25° (Bembenek et al. 2007; Richie et al. 2011). We utilized an ENU mutagenesis approach to isolate suppressors of *sep-1(e2406)* that result in viable F3 progeny at the restrictive temperature of 20° (see Materials and Methods) (Figure 1A). This approach yielded a total of sixty-eight independent suppressor lines from a total of 9.6 X 10^5^ haploid genomes (as determined by counting the approximate number of mutagenized F1 progeny). Each suppressor line was cloned and backcrossed with the original *sep-1(e2406)* line to reduce non-suppressing background mutations and homozygotes were isolated. A candidate gene sequencing approach was utilized to identify suppressor mutations within *sep-1*, *pph-5* and *hsp-90* (formerly known as *daf-21*) (see Materials and Methods). We also isolated six lines with novel unknown mutations belonging to at least four complementation groups.

### Intragenic suppressors of *sep-1(e2406)* are exclusively N-terminal

There were fourteen independent suppressor lines identified as intragenic *sep-1(e2406)* suppressors. All intragenic *sep-1(e2406)* suppressors resulted in missense mutations within the N-terminal region of SEP-1 and none are found in the catalytic domain of the protein (Figure 2A). Some mutations were identified from multiple independent lines. Interestingly, mutation in lysine 556 was identified in six lines.

**Figure 2.**
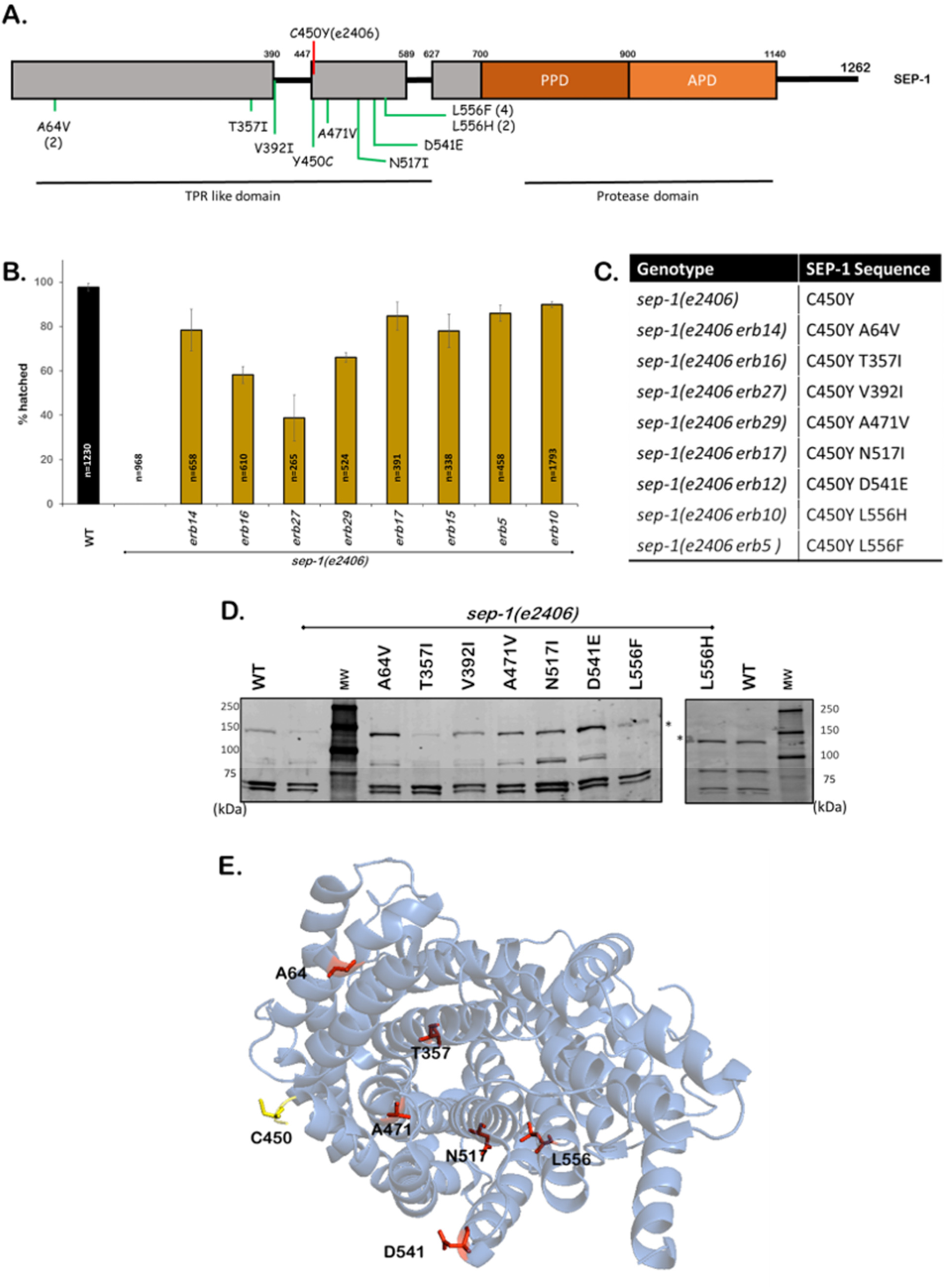
Intragenic mutations suppress *sep-1(e2406)* **A.** Protein diagram illustrating suppressing alleles of SEP-1. The causative *e2406* mutation (C450Y) is depicted with a red line. Novel suppressor mutations are in green on the protein diagram and are exclusively in the N-terminal TPR-like domain of the protein. Numbers in parentheses following a mutation indicate the number of times each suppressor was identified. **B.** Embryonic lethality assays demonstrate that each suppressing mutation restores viability to *sep-1(e2406)* worms at 20° (n=number of embryos). **C.** Table listing gene mutations and the resulting missense mutations in SEP-1. **D.** SEP-1 is detectable by western blot in animals carrying *sep-1(e2406)* suppressing mutations. Asterisk indicates SEP-1 (144KDa). **E.** Cryo-EM structure (PDB 5MZ6) illustrating the N-terminal TPR like domain of SEP-1. The residue mutated in *sep-1(e2406)* (C450) is shown in yellow and suppressor mutations are illustrated in red. Mapping of mutated residues onto the structure illustrates that they are distributed throughout the N-terminus.

The types of missense mutations observed include ones that increase the size of amino acid side chains while preserving charge (A64V, A392I, A471V and D541E). We also found mutations that remove charged side chains and introduce hydrophobic residues (T357I and N517I). The residue most frequently mutated was L556 and both changes we observed result in the introduction of aromatic side chains (L556F and L556H, Figure 2A, S2). It is also notable that L556F was previously identified as an intragenic *sep-1(e2406)* suppressor (Richie et al. 2011). We find that multiple residues in the N-terminus can be changed to restore function to the *sep-1(e2406)* mutant and restore viability (Figure 2 B and C).

One possible mechanism of suppression is that these mutations affect the stability of separase. To address this, we performed western blotting analysis of SEP-1 abundance in each of the suppressed lines. The separase protein is detectable in adult worms (Figure 2D) showing that proteins carrying suppressor mutations are expressed. Quantification shows that the original SEP-1(e2406) mutant protein is 40% as abundant as wild type SEP-1. The least effective rescuing mutation, *erb27* (V392I), is expressed at about twice the level of wild type separase. The three most effective rescuing mutations (*erb17* (N517I), *erb10* (L556H) and *erb5* (L556F)) have varying levels of expression. SEP-1(erb5) is 2.5-fold as abundant as wild type whereas SEP-1(erb17) and SEP-1(erb10) are expressed at 1.5-fold of wild type. The least abundantly expressed mutant, SEP-1(erb16) *(T357I)*, is not the least effective suppressor. No clear correlation is observed between protein abundance and rescuing ability, suggesting that these mutations do not simply affect protein levels, but may affect separase structure and function.

To gain more insight into these mutations, we mapped mutated suppressor residues onto the recently published Cryo-EM structure of SEP-1 in complex with its pseudosubstrate inhibitory chaperone; IFY-1 (securin) (PDB 5MZ6, Boland et al. 2017). This analysis reveals that there is no clustering of mutated residues to any specific surface in the TPR like domain of the N-terminus (Figure 2E). C450, the residue mutated in SEP-1(e2406), is at the edge of helix 16 and part of an unstructured loop containing about sixty amino acids between helix 15 and 16 of the TPR-like domain. The effects of C450Y mutation on the structure of the TPR-like domain have not been elucidated, but there is a potential that introducing a large aromatic residue on this solvent exposed loop may be unfavorable and could lead to a structural rearrangement of the SEP-1 N-terminal domain. The residues mutated in suppressed lines are found on helices not near C450, facing the interior of the protein and are likely involved in intramolecular interactions (Figure S2). These mutations have the potential of introducing new intramolecular interactions leading to improved structural stability of the SEP-1 TPR-like domain that may be disrupted in SEP-1(e2406). It is important to consider that the separase Cryo-EM structure represents a securin-bound fold of the enzyme, which is inactive. The active conformation of separase might bring these key residues into more obvious functionally relevant orientations.

### *pph-5* mutants are most frequently identified *sep-1(e2406)* suppressors

The majority of *sep-1(e2406)* suppressors identified from our analysis are mutations in the protein phosphatase, *pph-5*. The types of mutations identified include premature stop codons (12 alleles), splice site mutations (6 alleles), as well as amino acid substitutions (29 alleles). Missense *pph-5* suppressing mutations span the full length of the protein, altering both the TPR as well as the phosphatase domain. Excluding mutations that introduce a premature stop codon, our screen has identified twenty-five unique amino acid substitutions across the protein (Figure 3A). Missense suppressor mutations occur both within the TPR domain and the phosphatase domain of PPH-5, suggesting that both domains are required for PPH-5 regulation of separase. The capacity of these mutations to rescue *sep-1(e2406)* varies, as assayed by the proportion of embryos able to hatch at the restrictive temperature (Figure 3 B and C). Strong RNAi knockdown of *pph-5* (*pph-5 RNAi*) in these suppressed lines also results in improvement of suppression (Table 1) suggesting that they are reduction-of-function mutations.

**Table 1.** Reduction of *pph-5* by RNAi mediated knockdown results in improved hatching. RNAi knockdown of *pph-5* by feeding results in improved hatching efficiency in worms carrying *pph-5* mutations that suppress *sep-1(e2406)* lethality at the restrictive temperature of 20°.

| *sep-1* | *pph-5* | *pph-5* RNAi |  | No RNAi |
| --- | --- | --- | --- | --- |
|  |  | Total embryo | % hatching | % hatching |
| WT | WT | 512 | 98.1 | 97.6 |
| *e2406* | WT | 578 | 72.8 | 0.0 |
|  | *erb13* (Y65*) | 323 | 88.9 | 79.4 |
|  | *erb58* (I32N) | 255 | 98.0 | 73.3 |
|  | *erb47* (Y52H) | 235 | 73.5 | 70.4 |
|  | *erb51* (G66E) | 589 | 77.1 | 75.7 |
|  | *erb44* (L77P) | 226 | 95.1 | 60.2 |
|  | *erb54* (S105F) | 298 | 33.9 | 26.3 |
|  | *erb30* (M211K) | 113 | 94.7 | 92.9 |
|  | *erb46* (H243R) | 215 | 85.1 | 79.8 |
|  | *erb45* (G244E) | 604 | 78.3 | 78.9 |
|  | *erb31* (D270A) | 320 | 94.7 | 92.8 |
|  | *erb57* (D270N) | 434 | 97 | 73.4 |
|  | *erb48* (M285R) | 325 | 91.1 | 65.6 |
|  | *erb63* (R300C) | 322 | 75.5 | 39.6 |
|  | *erb21* (N309K) | 351 | 95.4 | 53.4 |
|  | *erb59* (M311R) | 378 | 65.5 | 58.6 |
|  | *erb42* (Y322I) | 270 | 79.3 | 51.1 |
|  | *erb72* (H351R) | 163 | 77.6 | 13.1 |
|  | *erb68* (S397P) | 292 | 27.3 | 28.2 |
|  | *erb33* (W413G) | 386 | 84.5 | 60.8 |
|  | *erb38* (C414Y) | 255 | 76.5 | 67.7 |
|  | *erb28* (H426Q) | 399 | 95.2 | 79.3 |
|  | *erb22* (C441Y) | 374 | 79.1 | 78.1 |
|  | *erb65* (T443I) | 399 | 84.4 | 57.7 |
|  | *erb40* (P448L) | 109 | 74.3 | 68.4 |
|  | *erb52* (G458E) | 282 | 78.0 | 61.3 |

**Figure 3.**
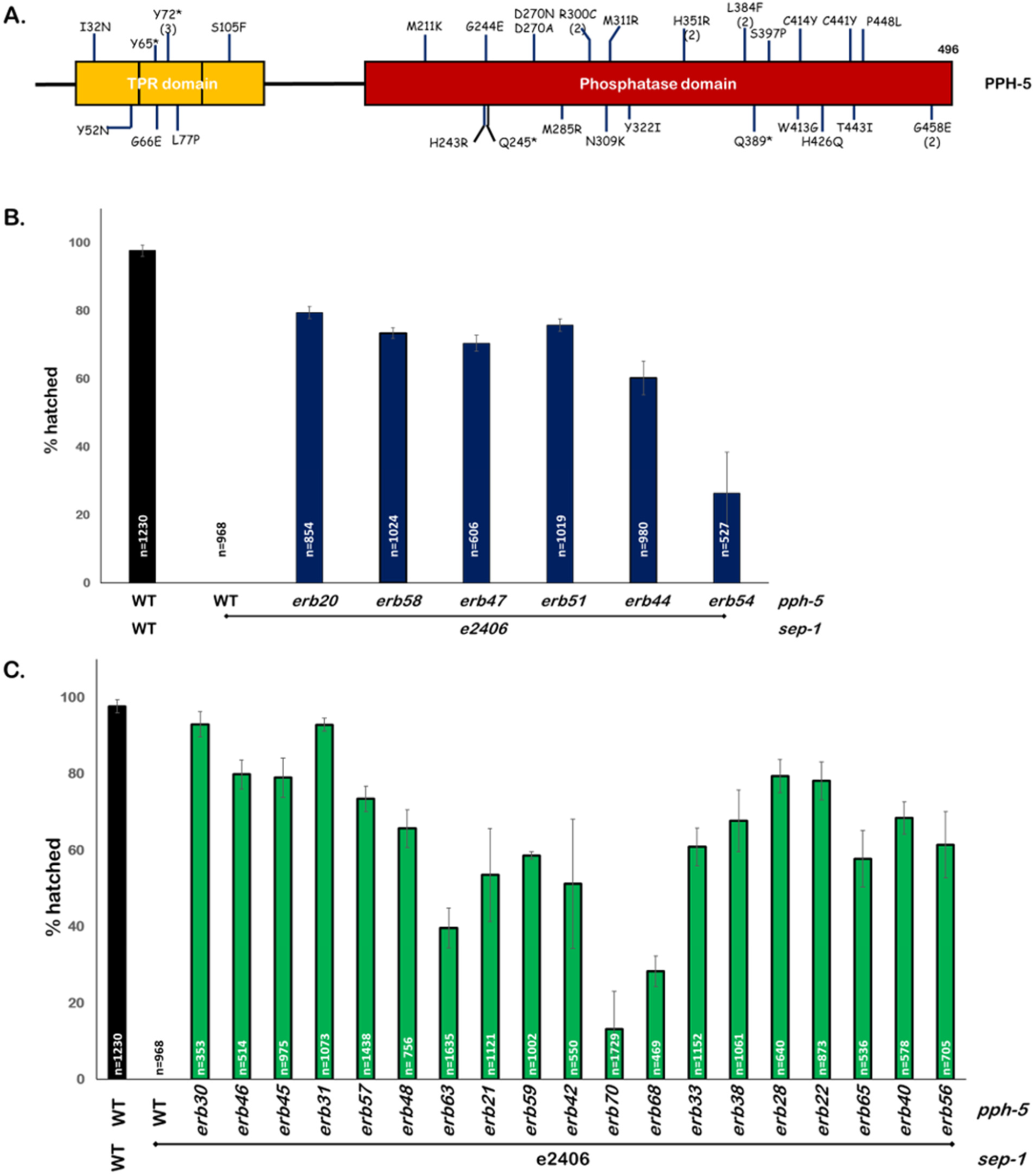
Mutations in *pph-5* are the most frequently identified *sep-1(e2406)* suppressors. **A.** Protein diagram illustrating *pph-5* alleles suppressing *sep-1(e2406)*. These missense mutations span across PPH-5. Numbers in parentheses following a mutation indicate the number of times each suppressor was identified. This diagram does not depict splice site variants or frameshift mutations. **B.** Embryonic lethality of *sep-1(e2406)* is rescued by missense mutations in the TPR domain of PPH-5, which might affect substrate binding. **C.** Embryonic lethality of *sep-1(e2406)* is rescued by missense mutations in the phosphatase domain of PPH-5, which might affect catalytic activity (n=number of embryos).

It has been shown that *pph-5* mutants do not suppress *sep-1(e2406)* by bypassing separase requirement (Richie et al. 2011) as RNAi knockdown of *sep-1* still results in lethality in suppressed lines. It is likely that the suppressors we have identified are *pph-5* reduction of function mutants and restore viability in a similar manner as previously identified *pph-5* mutants. Our data do not preclude the possibility that *pph-5* acts in a separase independent pathway to restore viability to *sep-1(e2406)* animals. We favor our proposed model because mutations in *pph-5* have been demonstrated to restore mutant separase localization (Richie et al. 2011). One suppressor mutation in *pph-5* (L77P) was independently identified in both screens as a suppressor of conditional (*sep-1(e2406)*) and non-conditional (*sep-1(ax110)*) separase mutants. This extensive collection of *pph-5* mutants provides a valuable tool for structure-function as well as genetic analysis of this phosphatase.

### HSP-90 suppressor reveals novel regulator of separase

The biochemical evidence connecting PPH-5 with HSP-90 (Haslbeck et al. 2015) prompted us to test if any of the suppressors were *hsp-90* mutations. We sequenced the *hsp-90* locus of the remaining suppressed lines that did not carry any suppressing intragenic or *pph-5* mutations. We found that *erb71* has a single missense mutation that changes methionine 661 into lysine (Figure 4A) that has an intermediate ability to restore hatching to 31% (Table 2). When isolated from *sep-1(e2406)*, the *hsp-90*(*erb71*) allele has minimal effect on embryonic survival at 20° (84% hatching) which suggests that the essential functions of HSP-90 are minimally affected (Figure 4B). The rescue observed with *pph-5(RNAi)* is greater than the 31% survival observed in *hsp-90*(*erb71)*. This suggests that either the M661L mutation does not completely disrupt the PPH-5 activating functions of HSP-90 or that PPH-5 can still be active without HSP-90. Consistent with this, we observed improved survival (92.9% hatching) when *pph-5(RNAi)* was performed in a *sep-1(e2406); hsp-90(erb71)* animal (Table 2). The identification of a HSP-90 allele that can suppress a temperature sensitive separase mutation is consistent with the hypothesis that HSP-90 acts via its regulation of PPH-5. Our data, however, do not exclude the possibility that HSP-90 directly regulates separase independent of PPH-5.

**Table 2.** RNAi mediated knockdown of *pph-5* in *hsp-90(erb71)* worms. The genetic interaction between *pph-5* and *hsp-90* was investigated by using RNAi mediated knockdown of *pph-5*. Reduction of PPH-5 in a worm carrying a *sep-1(e2406)* rescuing *hsp-90* mutation results in reduced embryonic lethality at the restrictive temperature (20°C). However, *pph-5(RNAi)* has little effect on embryonic lethality of *hsp-90(erb71)*.

| Strain | Total embryos | % hatching |
| --- | --- | --- |
| N2 | 512 | 98.4 |
| N2, *pph-5(RNAi)* | 352 | 97.8 |
| *sep-1(e2406)* | 223 | 0 |
| *sep-1(e2406); pph-5(RNAi)* | 340 | 71.1 |
| *sep-1(e2406); hsp-90(erb71)* | 181 | 30.9 |
| *sep-1(e2406); hsp-90(erb71); pph-5(RNAi)* | 157 | 92.9 |
| *hsp-90(erb71)* | 148 | 84.4 |
| *hsp-90(erb71); pph-5(RNAi)* | 186 | 86.5 |

**Figure 4.**
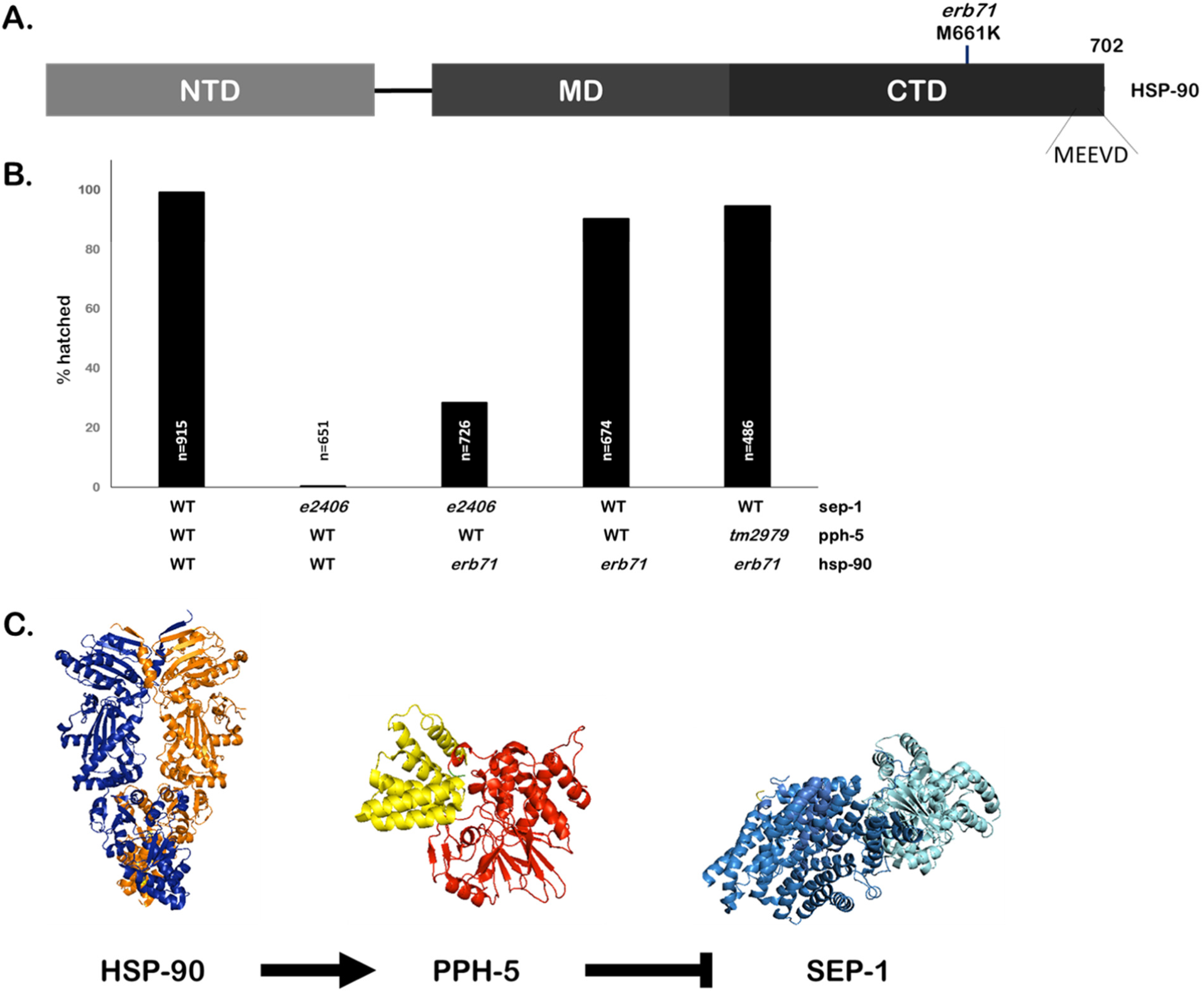
Mutation in the molecular chaperone *hsp-90* suppresses *sep-1(e2406)*. **A.** Protein diagram of HSP-90. The *erb71* mutation results in a missense mutation at the C-terminal end of the protein chaperone HSP-90 (M661K), separated by 36 residues from the C-terminal most MEEVD motif. HSP-90 protein domains are also illustrated (NTD, amino-terminal domain; MD, middle domain; CTD, carboxy-terminal domain; MEEVD, Met-Glu-Glu-Val-Asp motif) **B.** The *hsp-90(erb71)* mutant has minimal effect on hatching when present in an otherwise wild type background. Embryonic lethality is not reduced when *hsp-90(erb71)* is combined with a *pph-5* loss of function mutant (n=number of embryos). **C.** Model for separase regulatory pathway: HSP-90 activates PPH-5 to negatively regulate separase function. Loss of this negative regulation suppresses *sep-1(e2406)*. (PDB structures modified from 5MZ6 (SEP-1), 4JA9 (PPH-5) and 5FWP (HSP-90)).

Combining *hsp-90(erb71)* with *pph-5(RNAi)* has little effect on embryonic survival, compared to the effects of *erb71* alone (84% vs 86% hatching) in an otherwise wild type background. No significant changes in hatching were observed when *hsp-90(erb71)* was combined with *pph-5(tm2979)*. *pph-5(tm2979)* is an in-frame deletion that removes 55 amino acids from the PPH-5 TPR domain and potently suppresses *sep-1(e2406)* and *sep-1(ax110)* (Richie et al. 2011). These observations demonstrate that *pph-5* function is not critical, even in a mutant *hsp-90(erb71)* background, for the essential functions of HSP-90. Taken together, these observations support the hypothesis that *hsp-90(erb71)* does not result in a general loss in HSP-90 chaperone activity.

It is interesting to note that the mutation in HSP-90(erb71) (M661K) is found just N-terminal to the HSP-90 MEEVD motif, which is critical for HSP-90 to activate PPH-5 (Haslbeck et al. 2015). There is biochemical evidence that the PPH-5/HSP-90 interaction involves additional HSP-90 domains beyond the MEEVD motif. Activation of PPH-5 phosphatase by a peptide containing the MEEVD motif is less than that observed for full-length HSP-90 (Haslbeck et al. 2015). Crosslinking experiments also suggest additional contacts between HSP-90 and PPH-5. The corresponding residue mutated in *HSP-90(erb71)* in human HSP90 (M813) is part of the dimerization interface of two Hsp90 molecules near the site of TPR integrating MEEVD domain; as observed in a cryo-EM structure (PDB 5FWP, Verba et al. 2016). This residue might alter the ability of the MEEVD peptide to bind to the TPR domain of PPH-5 by altering HSP-90 C-terminal structure. Therefore, an analogous mutation in other organisms such as human cells may be useful for studies of the HSP90-PP5 pathway. We propose a model for the regulation of separase in which *pph-5* is a negative regulator of separase and PPH-5 activity is positively regulated by interactions with HSP-90 (Figure 4 C). These new alleles of *hsp-90* and *pph-5* provide important tools for future dissection of this pathway.

### Novel *sep-1(e2406)* suppressors belong to multiple complementation groups

Our suppressor screen identified six lines without suppressor mutations in the three genes we sequenced (*sep-1*, *pph-5* and *hsp-90*). These suppressed lines have varying degrees of hatching recovery at the restrictive temperature (Figure 5 A). To determine the number of loci represented by this group of alleles, we performed pairwise complementation tests. The hatching efficiency of broods laid by F1 cross progeny between two homozygous suppressed lines was monitored at 20°. As presented in Figure 5 B, these suppressors belong to four, possibly five, complementation groups. Two lines, *sep-1(e2406); erb23* and *sep-1(e2406)*; *erb24* do not complement and their cross progeny demonstrate an intermediate embryonic lethality as compared to the parents. Another mutant, *sep-1(e2406); erb66* mutation appears to be dominant over other suppressors, except *erb67*, and cannot be assigned to a complementation group. Finally, *erb37*, *erb60* and *erb67* do not result in suppression when crossed with other mutants and are likely mutations in three different genes. These observations provide an exciting opportunity to identify novel regulators of separase.

**Figure 5.**
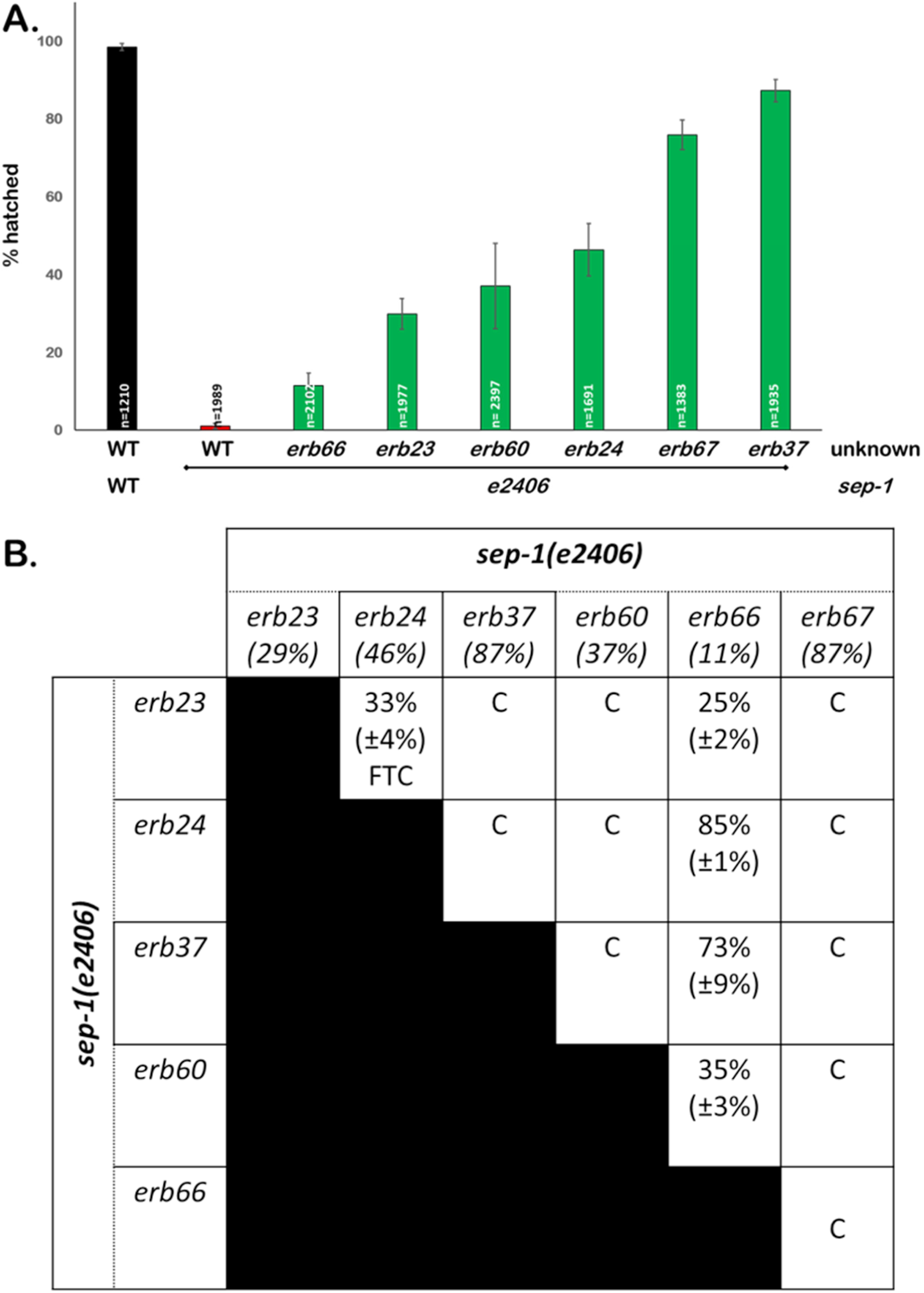
Novel suppressors of *sep-1(e2406)* belong to multiple complementation groups. **A.** Strains carrying novel *sep-1(e2406)* suppressors result in varied rescue of embryonic lethality (n=number of embryos). **B.** Complementation assay based on survival of F2 embryos of a cross between strains carrying novel *sep-1(e2406)* suppressors indicates these suppressors belong to multiple complementation groups. Numbers below each parent strain or in a box representing a cross progeny indicate the percent of embryos that hatch at the restrictive temperature of 20°. The numbers in parenthesis are standard deviations for three replicate hatching assays; C = Complements, FTC = Failure To Complement.

## Conclusion

By undertaking this extensive suppressor screen, we set out to identify separase regulators. Our results reveal that the phosphatase, *pph-5*, is a suppressor of *sep-1(e2406)* lethality. The results of our genetic screen highlight the importance of the *pph-5* regulatory pathway. The mechanism by which *pph-5* regulates separase during cytokinesis will be an important focus of future studies. Identifying substrates of PPH-5 that become hyperphosphorylated in a *pph-5* mutant may elucidate this mechanism as well as any additional roles PPH-5 might play during mitosis. We have found that *hsp-90* also functions, likely via its regulation of *pph-5*, as a separase regulator. The sole *hsp-90* suppressor we identified may be a rare hypomorphic mutant whose PPH-5 activating role is selectively reduced without compromising its other critical chaperone functions. Given the high degree of conservation of *pph-5* and *hsp-90*, we expect our observations will be applicable to separase function in other systems as well. We were also able to identify novel intragenic suppressors, all of which are missense mutations in the N-terminal TPR-like domain of SEP-1, providing insight into this poorly characterized domain. TPR domains mediate protein-protein interactions and these residues may be involved in mediating interactions with separase binding partners required for its function. We have additionally isolated lines that carry mutations belonging to at least four complementation groups, giving us the opportunity to more extensively understand separase regulation. We will pursue a whole genome sequencing approach to identify these mutations. This study demonstrates the power of genetics in understanding separase function and regulation.

## ACKNOWLEDGMENTS

We would like to thank Aude Peden for performing the ENU screen and isolating suppressors. We would like to thank Chris Turpin for helping with the screen and discussion of the manuscript. We would like to thank Andy Golden from the National Institute of Diabetes & Digestive & Kidney Disease at the National Institutes of Health for his scientific discussion and guidance. We would also like to thank Kevin O'Connell and Bruce McKee for providing critical feedback on the manuscript. We especially thank Wormbase and the Caenorhabditis Genetics Center (CGC). WormBase is supported by grant U41 HG002223 from the National Human Genome Research Institute at the US National Institutes of Health, the UK Medical Research Council and the UK Biotechnology and Biological Sciences Research Council. The CGC (St. Paul, MN), is funded by the National Institutes of Health Office of Research Infrastructure Programs (P40 OD010440). This work was funded by startup funds from UT Knoxville and by NIH R01 GM114471.

